# iFORM: incorporating Find Occurrence of Regulatory Motifs

**DOI:** 10.1101/044214

**Authors:** Chao Ren, Hebing Chen, Feng Liu, Hao Li, Xiaochen Bo, Wenjie Shu

## Abstract

**Motivation:** Accurately identifying binding sites of transcription factors (TFs) is crucial to understand the mechanisms of transcriptional regulation and human disease.

**Results:** We present incorporating Find Occurrence of Regulatory Motifs (iFORM), an easy-to-use tool for scanning DNA sequence with TF motifs described as position weight matrices (PWMs). iFORM achieves higher accuracy and sensitivity by integrating the results from five classical motif discovery programs based on Fisher’s combined probability test. We have used iFORM to provide accurate results on a variety of data in the ENCODE Project and the NIH Roadmap Epigenomics Project, and has demonstrated its utility to further understand individual roles of functional elements.

**Availability:** iFORM can be freely accessed athttps://github.com/wenjiegroup/iFORM.

## Introduction

Gene regulation is coordinately regulated by interactions of many transcription factors (TFs), many of which bind DNA in the promoters and enhancers preferentially at characteristic sequence ‘motif’ that is a short pattern and tends to be conserved by purifying selection. Identifying and understanding of these motifs can provide critical insight into the mechanisms of transcriptional regulation and human disease. However, accurate identification of these motif binding sites is still challenging because a single TF will often recognize a variety of similar sequences.

Over the past several decades, many classical algorithms have been developed to discovery DNA regulatory motifs. FIMO (Grant, et al., 2011), Consensus(Hertz and Stormo, 1999; Stormo and Hartzell, 1989), and STORM (Schones, et al., 2007) have unitary function of motif scanner, while RSAT (Thomas-Chollier, et al., 2011) and HOMER (Heinz, et al., 2010) provide multiple functions in analysing regulatory sequences in general. Table S1 summarized the features of iFORM and these five algorithms as motif scanner. Each of these methods has its own merits in identifying potential TF binding, however, it is still a critical challenge to integrate superiorities and to preclude inferiorities of these complementary methods. Several studies have demonstrated that higher accuracy and sensitivity can be achieved by incorporating multiple motif discovery programs (Harbison, et al., 2004; MacIsaac and Fraenkel, 2006; Tompa, et al., 2005), however, this will result in considerable computational overhead for scoring the huge number of discovered motif instances obtained with different programs.

We describe incorporating Find Occurrence of Regulatory Motifs (iFORM), a software tool for scanning DNA sequence with TF motifs described as position weight matrices (PWMs) by integrating five classical motif discovery methods. We used Fisher’s combined probability test to convert the resulted *p*-values obtained from the five methods to a *χ*^2^ statistic, which follows a chi-squared distribution. We then applies false discovery rate analysis to estimate a *q*-value for each motif instance. By systematically assessing the accuracy of prediction performance, iFORM achieve higher accuracy and sensitivity relative to five classical algorithms. Although iFORM integrated five methods, it is efficient, allowing for scanning sequences at a rate of 3.5Mb/s on a single CPU. Additionally, we have illustrated the use of iFORM by producing high-quality genome-wide maps of the TFBSs for 542 TFs within DHSs of 133 human cell and tissue types that were generated in the ENCODE Project (Consortium, 2012) and the NIH Roadmap Epigenomics Mapping Consortium (Bernstein, et al., 2010) in our recent studies. We also demonstrated the utility of iFORM to further understand individual roles of functional elements in gene regulation and disease based on these maps.

## Results

### Implementation of iFORM

iFORM integrated the motif instances identified by five classical algorithms including FIMO (Grant, et al., 2011), Consensus(Hertz and Stormo, 1999; Stormo and Hartzell, 1989), STORM (Schones, et al., 2007), RSAT (Thomas-Chollier, et al., 2011) and HOMER (Heinz, et al., 2010) based on Fisher’s method. iFORM was built using C, Perl, and Python, and the overview of workflow is illustrated in Fig. S1. To improve the efficiency, we extracted the core source code of the function of motif discovery from these five algorithms and integrated them in the framework of iFORM, instead of just combining the resulted *p*-values obtained from these algorithms.

iFORM accepts DNS sequences in FASTA format or genomic coordinates in BED or GFF formats, and takes as input one or more TF motifs (represented as PWMs) that can be collected form an existing motif database or generated from the MEME algorithm, or even defined by user self. For each motif instance, iFORM computes a *χ*^2^ statistic that obtained by combining *p*-values resulted from the five algorithms using Fisher's method, and converts these statistics to *p*-values using chi-squared distribution. Finally, iFORM used a bootstrap method (Storey, 2002) to estimate false discovery rates (FDRs), and reported for each *p*-value a corresponding *q*-value, which is defined as the minimal FDR threshold at which the *p*-value is deemed significant (Storey, 2003).

iFORM outputs a ranked list of motif instances, each with an associated *χ*^2^ statistic, *p*-value and *q*-value. The list is illustrated as an HTML report, as an XML file in CisML format (Haverty and Weng, 2004), as a plain text file and as tab-delimited files in gff and wig formats suitable for input to the UCSC Genome Browser (Speir, et al., 2016).

### Performance assessment of iFORM

Since iFORM integrates five classical algorithms based on Fisher’s method, it is expected that iFORM can achieve higher accuracy and sensitivity comparing with the five methods. To test it, we first examined the CTCF motif instances that are identified by these six methods separately. We found that many of the CTCF binding sites, which are well annotated by DNase-seq, DGF, and corresponding TF ChIP-Seq data in H1 cells, can only be discovered by iFORM, and cannot be detected by other five classical methods (Fig. S2A). To systematically assess the accuracy of prediction performance for each motif instance, we used receiver operation curves (ROCs) and corresponding the area under the curve (AUC) on the “gold-standard” data of six TF ChIP-seq data in GM12878 cell provided in a previous study (Pique-Regi, et al., 2011) (Fig. 1A and Fig. S). Our results suggest that our method iFORM showed both higher sensitivity and specificity than other five classical method.

**Figure 1.**
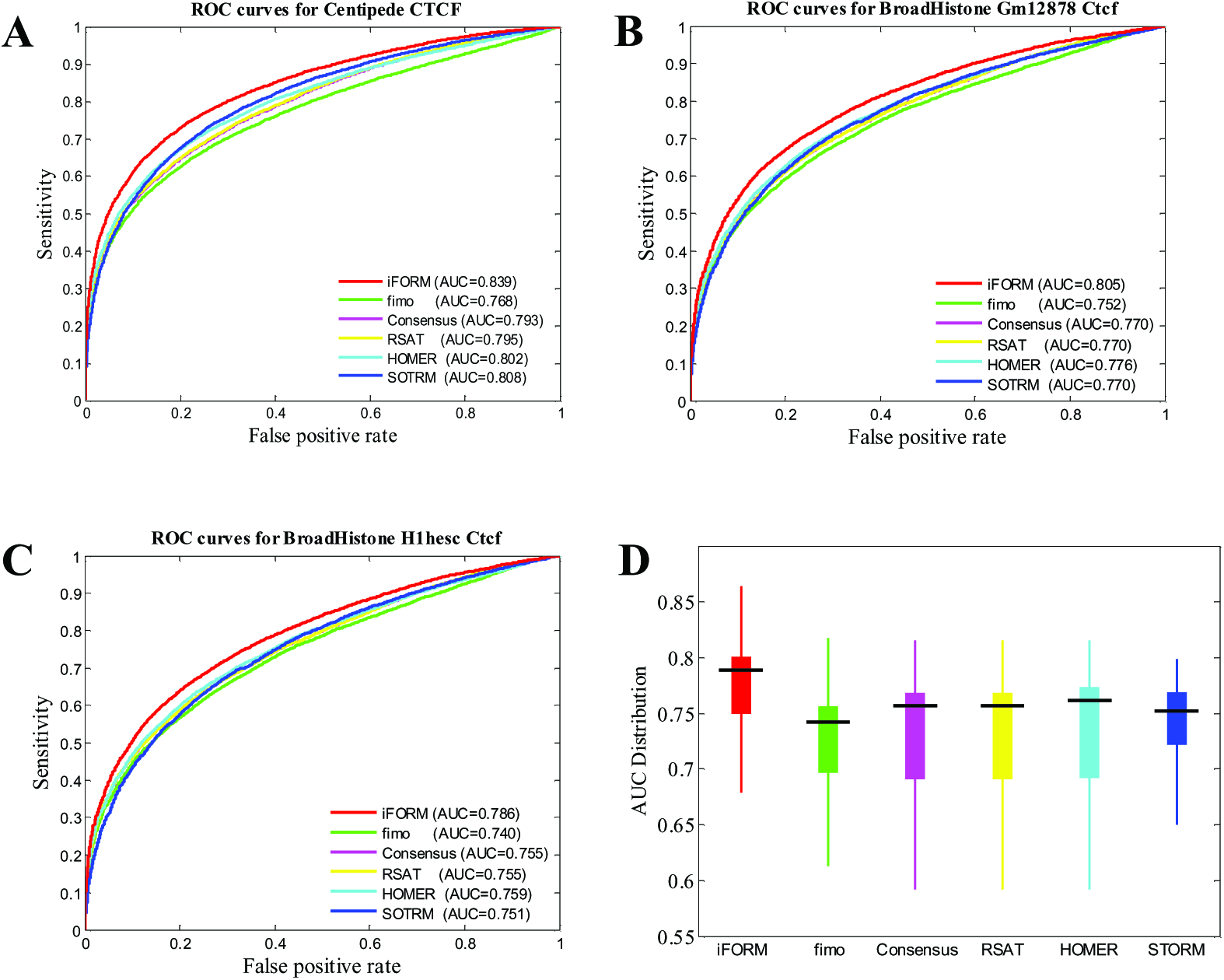
Performance assessment of iFORM. (A) ROC curves of multiple algorithms using the “gold-standard” data of CTCF in GM12878 cells provided in a previous study. (B) ROC curves of multiple algorithms using the “gold-standard” data of CTCF in GM12878 cells that was generated in our study. (C) ROC curves of multiple algorithms using the “gold-standard” data of CTCF from H1 cells. (D) Boxplot distribution of AUC for diverse TFs in multiple cells/tissues.

To further validate the predicated TF binding sites, we used the method presented by Roger Pique-Regi *et al* (Pique-Regi, et al., 2011) to generate the “gold-standard” data for many TF ChIP-seq that were produced from different laboratories and from different cells/tissues. Using our newly generated “gold-standard” data of CTCF in GM12878 cell, we obtained similar ROCs as presented in Fig. 1A (Fig. 2B). In addition, similar ROCs also obtained for the newly generated “gold-standard” data of CTCF in GM12878 produced from different labs (Fig. S2B-D), and for the newly generated “gold-standard” data of CTCF in other cells (Fig.S2E). Across these “gold-standard” data, our iFORM illustrates higher AUCs than other five algorithms. These results suggest that iFORM manifested superior performance consistency in different laboratories and different cells/tissues and demonstrated state-of-the-art performance relative to five existing methods.

Next, we assessed the prediction performance using new “gold-standard” data of other 109 TF ChIP-seq data produced in ENCODE project (Table S3). iFORM illustrates significant higher AUC than those of the five classical methods (Fig. 2D, *p*-value < 10^−6^). Taken together, these results indicate that our novel method iFORM achieve higher accuracy and sensitivity comparing with the five classical methods.

### Identification of TFBSs with iFORM

We produced high-quality genome-wide maps of the TFBSs for 542 TFs within DHSs of 133 human cell and tissue types that were generated in the ENCODE Project (Consortium, 2012) and the NIH Roadmap Epigenomics Mapping Consortium (Bernstein, et al., 2010) using iFORM (*p*-value < 10^−18^). On average, we obtained approximately 4,470 TFBSs for each TF and for each cell/tissue.

Based on these data, we found that TFBSs were clustered together in the human genome, and report the first comprehensive map of TFBS-clustered regions across human cell and tissue types. An integrative analysis of these regions revealed a novel transcriptional regulation model on the accessible chromatin landscape (Chen, et al., 2015). Additionally, we investigated the HOT (high-occupancy target) regions, which were defined as TFBS-clustered regions with extremely high TFBS complexity. We found that HOT regions play key roles in human cell development and differentiation (Li, et al., 2016). Furthermore, we explored the association of GWAS SNPs and HOT regions, and demonstrate the key roles of HOT regions in human disease and cancer (Li, et al., 2015). These findings represent a critical step toward further understanding disease biology, diagnosis, and therapy.

## Conclusions

The iFORM presented in this study was designed to incorporate five classical regulatory motif discovery methods using Fisher’s method. iFORM is an easy-to-use motif discovery tool that achieve higher accuracy and sensitivity by integrating the results from multiple motif discovery programs. The iFORM has provided accurate results on a variety of data in the ENCODE Project and the NIH Roadmap Epigenomics Project in our recent studies, and has been demonstrated its utility to further understand individual roles of functional elements in the mechanisms of transcriptional regulation and human disease.

## Materials and Methods

### Data sets

DNaseI Hypersensitivity by Digital DNaseI data were obtained from both the ENCODE Project (Consortium, 2012) and the NIH Roadmap Epigenomics Mapping Consortium (Bernstein, et al., 2010). Transcription factors by ChIP-seq data were obtained from the ENCODE Project (Consortium, 2012). The uniform processing pipeline of ENCODE Integrative Analysis Consortium was used to generate the uniform peaks of both DNase-seq and ChIP-seq data. The use of these data strictly adheres to the ENCODE and Roadmap Epigenomics Consortium Data Release Policy. PhastCons were extracted from the hg19 conservation track of the UCSC Genome Browser (Speir, et al., 2016).

The position weight matrices (PWMs) of the 542 TFs, corresponding to 796 motif models, were collected from the TRANSFAC (Matys, et al., 2006), JASPAR (Portales-Casamar, et al., 2010), and UniPROBE (Robasky and Bulyk, 2011) databases as described in our previous study.

### Incorporating find occurrence of regulatory motifs

iFORM integrates the TF binding sites identified by five classical method including FIMO (Grant, et al., 2011), Consensus(Hertz and Stormo, 1999; Stormo and Hartzell, 1989), STORM (Schones, et al., 2007), RSAT (Thomas-Chollier, et al., 2011) and HOMER (Heinz, et al., 2010). For each discovered TFBS, the combined *p*-values obtained from these five methods was calculated using Fisher's combined probability test. First, a test statistic is calculated using the formula:

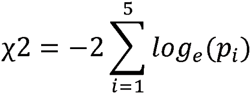

where *p*_*i*_ are the *p*-values calculated from the five methods for each TFBS. *χ*^2^ follows a chi-squared distribution *χ*^2^(2*5), thus a combined *p*-value can be assigned to this test statistic. A *p*-value threshold of enrichment of 10^−9^ was used for all data sets.

### Generation of the “gold-standard” data

To validate the predicted TFBSs using ROC approach, we applied a similar method presented in a previous study (Pique-Regi, et al., 2011) to generate “gold standard” data for a few TFs. Briefly, we scanned the genomic sequences under the uniform DHSs in the hg19 genome using the five methods incorporated by iFORM with default parameters for each TF motif in the set of “gold standard”, separately. Then, for each TF, all motif instance (*p*-value < 10^−5^) that located within the corresponding TF ChIP-seq peaks were considered as the set of TFBS positives. The set of TFBS negatives were defined as all motif instance (*p*-value < 10^−5^) that did not overlap a ChIP-seq peaks and had a greater fraction of total mapped reads from the “Control” as compared to the ChIP-seq treatment. Totally, 109 sets of “gold standard” for 12 TFs from 51 cell/tissues (Table S2).

### Assessing the performance of iFORM

The prediction performance of iFORM was assessed using three methods, including a correlation based approach, Receiver Operation Curves (ROCs), and conservation. First, we used Receiver operation curves (ROCs) and the area under the curve (AUC) to assess the accuracy of prediction performance of iFORM.

In contrast to ROC analysis that examined the motif instances that are either TFBS positives or TFBS negatives, we additionally adopted a correlation approach that takes into account almost all locations, except those in or near repetitive regions. The correlation approach assumes that the larger the correlation between the predicted values and the ChIP-seq signal and the smaller the correlation with the background noise “Control”, the better is the prediction accuracy. We extracted the ChIP-seq reads and the “Control” reads around each motif site, and used the Pearson correlation to measure the trend between the square root of the total number of reads and the posterior log-odds reported by iFORM and the five classical methods.

For motifs where ChIP-seq data were not available, we used sequence conservation to assess whether iFORM was correctly detecting TF binding. For this, we withheld the phastCons score when fitting our model and defined a test statistic (conservation Z-score) that measured the significance of the logistic regression of the phastCons score of the motif on the posterior probability of binding (for full details, see Supplemental material).

## Acknowledgements

We wish to thank the ENCODE Project Consortium and the Roadmap Epigenomics Project Consortium for making their data publicly available. This work was supported by grants from the Major Research plan of the National Natural Science Foundation of China (No. U1435222), the Program of International S&T Cooperation (No. 2014DFB30020) and the National High Technology Research and Development Program of China (No. 2015AA020108).

## Author Contributions

W.S.: conceived the project.
W.S.: and
X.B.: designed all experiments.
C.R.: and
H.C.: wrote the program.
H.L.: and
F.L.: performed the data analysis.
W.S.: wrote the manuscript.

## Additional information

Supplementary information accompanies this paper at http://www.nature.com/scientificreports.

Competing financial interests: The authors declare no competing financial interests.

## Additional files

**Figure S1. The workflow of iFORM**

The overview of iFORM workflow. (1) Assemble input data. Results may be improved by restricting the input to high-confidence sequences. Some programs achieve improved performance by using phylogenetic conservation information from orthologous sequences or information about protein DNA-binding domains. (2) Choose several motif discovery programs for the analysis. For recommended programs see Figure 3. (3) Test the statistical significance of the resulting motifs. Use control calculations to estimate the empirical distribution of scores produced by each program on random data. (4) Clustering and post-processing the motifs. Motif discovery analyses often produce many similar motifs, which may be combined using clustering. Phylogenetic conservation information may be used to filter out statistically significant, but non-conserved motifs that are more likely to correspond to spurious sequence patterns. (5) Interpretation of motifs. Algorithms exist for linking motifs to transcript factors and for combining motif discovery with expression data.

**Figure S2. Performance comparisons between iFORM and existing methods**

An example of CTCF motif binding site, which are well annotated by DNase-seq, DGF, and corresponding TF ChIP-Seq data in H1 cells, can only be discovered by iFORM, but not by other five classical methods.

**Figure S3. ROC curves of CTCF in GM12878 cells from multiple labs**

ROC curves of multiple algorithms using the “gold-standard” data of CTCF in GM12878 cell that was generated from different labs.

**Figure S4. ROC curves of CTCF in diverse cells**

ROC curves of multiple algorithms using the “gold-standard” data of CTCF from diverse human cells/tissues.

**Figure S5. ROC curves of many TFs across diverse cells/tissues**

**Table legend**

**Table S1. Summaries of five motif scanners.**

**Table S2. Summary of “gold-standard” data.**

**Table S3. AUC of different TFs, related to Figure 1D.**

